# LiF-MS: Mapping unstructured peptide-protein interactions using Ligand-Footprinting Mass Spectrometry

**DOI:** 10.1101/361857

**Authors:** Benjamin Parker, Edward Goncz, David T. Krist, Alexander Statsyuk, Alexey I. Nesvizhskii, Eric Weiss

## Abstract

Unstructured peptides, or linear motifs, present a poorly understood molecular language within the context of cellular signaling. These modular regions are often short, unstructured and interact weakly and transiently with folded target proteins. Thus, they are difficult to study with conventional structural biology methods. We present Ligand-Footprinting Mass Spectrometry, or LiF-MS, as a method of mapping the binding sites and dynamic disorder of these peptides on folded protein domains. LiF-MS uses a cleavable crosslinker to mark regions of a protein contacted by a bound linear motif. We demonstrate this method can detect both conformation ensembles and binding orientations of a linear motif in its binding pocket to amino-acid-level detail. Furthermore, marked amino acids can be used as constraints in peptide-protein docking simulations to improve model quality. In conclusion, LiF-MS proves a simple and novel method of elucidating peptide docking structural data not accessible by other methods in the context of a purified system.

## Introduction

Short linear interaction motifs (SLiMs) are short, unstructured peptides in a protein which often mediate transient interactions with structured protein domains and function to fine-tune cellular process such as transcription and signaling. It is estimated there may be close to a million unique linear motifs in the human proteome^1^, many of which are hypothesized to be underrepresented in connections to disease^2^. Linear motifs interact with binding surfaces through divergent and difficult to predict interaction conformations and represent a hidden and poorly understood molecular language distinct from that of folded domain-domain interactions. Notably, while SLiM binding interactions in SH2 domains^3,4^ and MAP kinase (MAPK) D-sites^5,6^ are well studied, many hundreds of SLiM peptide families remain poorly understood. Furthermore, structural studies often rely on time-consuming techniques such as X-ray crystallography and NMR which cannot easily account for the intrinsic disorder often present in linear motif-binding domain interactions^5,7^.

Here, we present Ligand-Footprinting Mass Spectrometry (LiF-MS), a novel method for mapping both the binding footprint and structural dynamics of disordered peptide motifs within a folded binding pocket. While broadly applicable to the general study of intrinsically disordered protein-protein interactions, we demonstrate the power of LiF-MS by mapping the docking site and binding dynamics of the MKK4 docking motif, or D-motif, onto the human MAP kinases JNK1, ERK2, and p38α. The MKK4 D-motif is unusually non-specific in its binding and is the only known D-motif to bind the structurally divergent clefts of the JNK and ERK/p38 families^5,6,8–10^. While no crystal structure of MKK4 in complex with a kinase has been solved, its D-motif is hypothesized to form multiple, distinct MAPK-dependent conformations^5,6^.

Many previous studies on structural mapping utilize crosslinking reagents that covalently connect proximal protein regions, creating a variety of irreversibly linked tryptic peptides. Processing data from these experiments can be challenging as the resulting linked peptides can produce difficult to decrypt MS/MS spectra. As an alternative, cleavable crosslinkers which leave residual mass labels can be utilized to study disordered proteins. LiF-MS utilizes a diazirine-based photocrosslinker described by Krist et al.^11^ which can be cleaved by acid to leave a butanol (+72 dalton) mass mark. While not previously tested on disordered proteins, this crosslinker has been used to successfully map interactions of ubiquitin ligases with their cognate E2 and E2~Ub thioester mimics^11^.

By using MKK4-MAPK interactions as a platform, we demonstrate the capability of LiF-MS to map an ensemble of peptide conformations within a folded protein domain. We show that the MKK4 D-motif binds the common docking (CD) groove of MAPKs JNK1 and p38α in distinct conformations, confirming previous hypotheses^5,6^. We further show this peptide uniquely binds in a bi-directional manner to the kinase ERK2. Thus, we demonstrate how this methodology can be used to map the motion of an intrinsically disordered peptide within a folded protein domain.

## Results

### Ligand-Footprinting Mass Spectrometry (LiF-MS) methodology

Current methodologies to identify binding sites of small peptides on a folded protein domain require difficult to obtain crystallography or NMR data. We therefore sought to develop a simple and robust method for mapping SLiM interaction regions on a folded protein domain. We took advantage of a previously established photoactivatable crosslinker. This crosslinker contains: (1), a diazirine moiety which forms a reactive carbene upon irradiation with ultraviolet light, (2), a sulfur-reactive iodoacetamide for attachment to a site-engineered cysteine residue, and (3), an acid-cleavable sulfamide which leaves a butanol mark on the target (Fig. 1a). Diazirine crosslinking is non-specific, occurring with N-H, C-H, O-H, and other chemical bonds both in macromolecules and in solvent^12,13^. As a result, the majority of carbene is quenched by solvent and total crosslinking yield is lower than other chemistries. However, the higher stringency compensates for off-target crosslinking caused by the binding promiscuity of SLiMs and only detects specific protein-peptide interactions^12^. These elements make diazirines ideal for screening the binding sites of small peptides.

**Figure 1.**
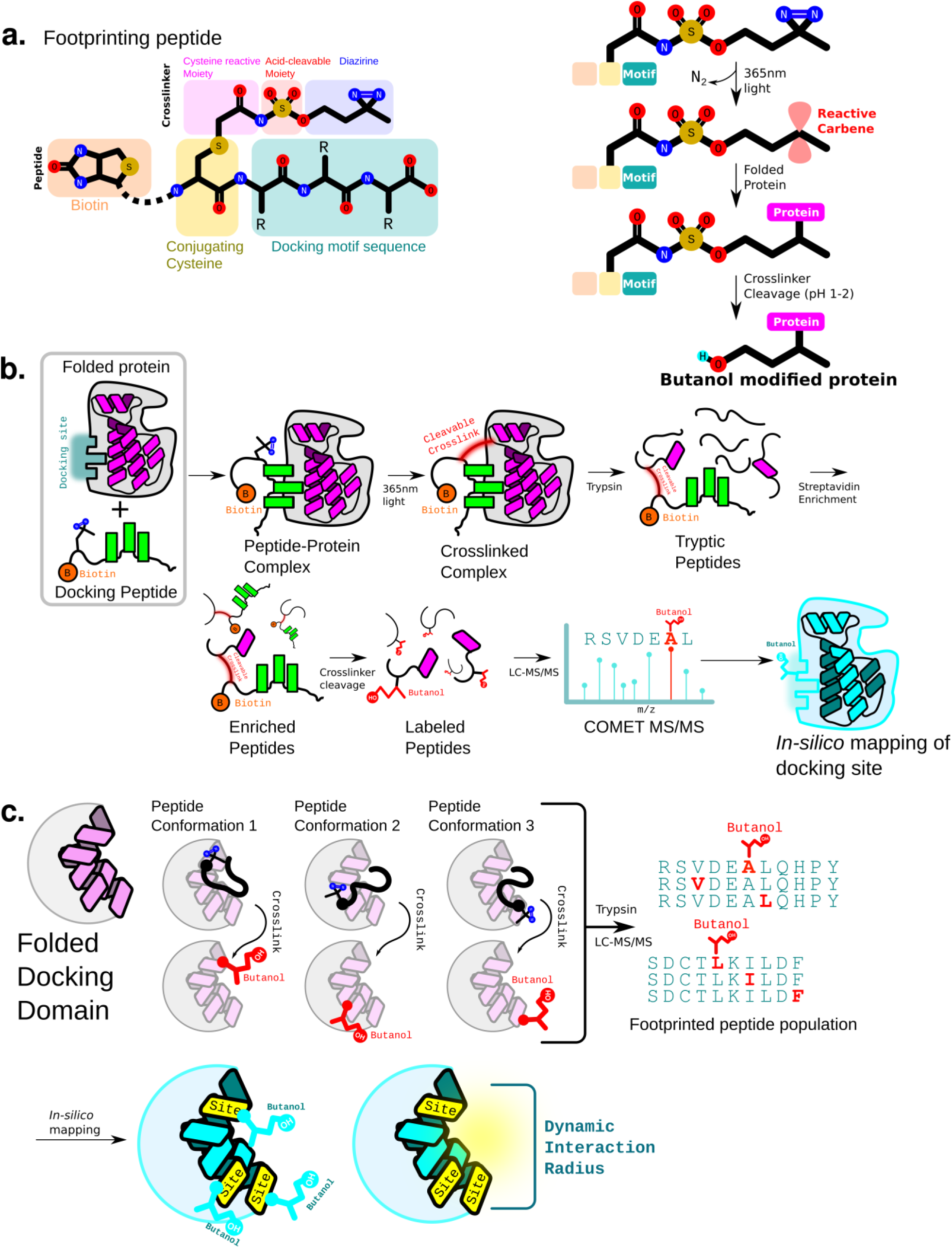
Overview of the LiF-MS workflow. (a) General structure of synthetic docking peptides used in this study as well as the diazirine-based crosslinker. Peptides are alkylated with an SN2 reaction between the iodoacetamide group and an added cysteine within the peptide. Irreversible crosslinking followed by cleavage yields a 72 dalton butanol mass mark. (b) LiF-MS footprinting methodology. Biotinylated peptides are crosslinked to folded protein domains, the crosslinker cleaved, and LC-MS/MS used to detect peptide docking sites. The COMET application is used to process proteomics data. (c) LiF-MS can detect conformational dynamics exhibited by disordered docking peptides in their binding pocket. Each population of peptide conformations is detected by LC-MS/MS and can be used to map a radius of interaction.

To map binding sites, we attached the crosslinker to a synthetic, biotinylated peptide containing the SLiM of interest (Fig. 1a). Incubation of this peptide with the protein of interest, followed by ultraviolet irradiation, results in a population of target protein covalently bound to the peptide at the interaction site. Trypsinization of the conjugated complex is followed by enrichment of crosslinked tryptic peptides with streptavidin agarose. Elution from the resin by cleavage of the crosslinker at low pH yields a solution of tryptic peptides covalently modified by a butanol mass tag. This is then used to map the motif’s binding site by LC-MS (Fig. 1b). Each crosslinked residue in the folded protein is scored based on detected mass shifts in Peptide Spectral Matches (PSMs) (Supplementary Fig. 1; also see Methods), and the resulting residues are mapped to the protein *in silico* (Fig 1b).

Distinct from folded protein-protein interfaces, motif-target interactions generally have high conformational heterogeneity^14^. This “fuzziness” is characteristic of intrinsically disordered regions^15^. LiF-MS hence generates an ensemble of protein-motif crosslinking events representing the dynamic motion of the motif in the binding pocket (Fig. 1c). Once mapped back onto the folded domain, it elucidates both the location of the docking site and the effective range of peptide conformations.

### Crosslinking synthetic D-motifs to purified MAP kinases

To test the utility of LiF-MS, we used the well-characterized MAPK to D-motif interaction as a platform. We chose to examine the interaction of a synthetic MKK4 D-motif to the human MAPKs JNK1, ERK2, and p38α since this interaction is relatively well characterized biochemically and behaves as a canonical D-motif^5,8,9,16^. Interestingly, MKK4 uniquely docks with all three MAPKs and is hypothesized to adopt an MKK6-like conformation when in complex with ERK and p38α kinases and an NFAT4-like conformation when bound to JNK1^5^. MKK6 and NFAT4 interact with the acidic CD-groove of the kinase in different ways. While NFAT4 forms a more linear structure, MKK6 is hypothesized to form a loop within the wider CD-groove of p38α^5,6^. A crystal structure of MKK4 in complex with a MAPK has yet to be solved, thus we hypothesized our methodology might also shed light on the unusual binding mechanism of this D-motif.

We synthesized a series of MKK4 D-motifs (referred to broadly as MKK4tides) based on previous work (Fig. 2a)^8^. Peptides contained an N-terminal biotin which we used for enrichment and crosslinking detection via western blot and a cysteine for crosslinker attachment. Using a minimal wild-type peptide, we found crosslinking to JNK1 occurred in the presence but not the absence of UV (Fig. 2b). However, as this peptide in practice yielded a weak signal in mass spectrometry experiments, we designed two additional shorter peptides (peptide I and II) (Fig. 2a) that cross-linked effectively to all three purified MAPKs (Fig. 2b,c).

**Figure 2.**
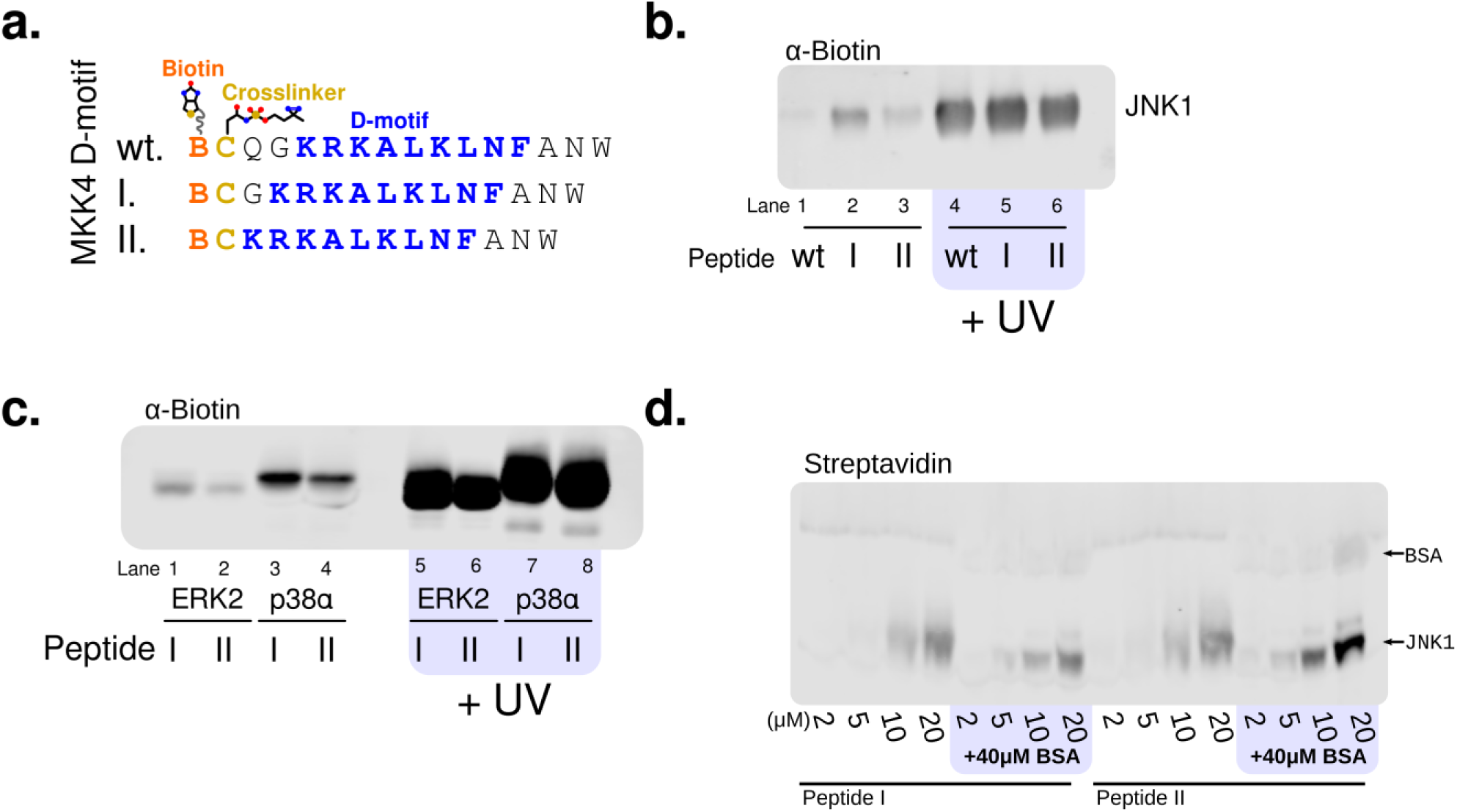
D-motifs in this study crosslink to MAP kinases. (a) Sequence of peptides used in this study. Full-length “wild type” peptide is QGKRKALKLNF. Yellow, (added) cysteines used to conjugate the crosslinker. Orange (B), biotin. (b-d) western blots of purified MAP kinases crosslinked to indicated peptides. Blots (b,c) were probed with anti-biotin antibody; blot (d) was probed with IRDye^®^ 800CW streptavidin (LI-COR). (b) Wild-type and truncated peptides I and II crosslink to purified JNK1 in a UV-dependent manner. (c) Peptides I and II exhibit UV-dependent crosslinking to purified ERK2 and p38α. (d) Concentration-dependent crosslinking of peptides I and II to JNK1. Samples containing 10μM JNK1 along with the indicated amounts of crosslinker-containing MKK4 D-motif peptide were irradiated with UV in the absence or presence of 40μM BSA.

Diazirines are reported to have low non-specific crosslinking^12^. We expect the high stringency of the diazirine warhead to only efficiently crosslink when bound in a specific conformation to a target protein and not when interacting non-specifically. To confirm this, we crosslinked increasing concentrations of peptides I and II to purified JNK1 in a molar excess of BSA (Fig. 2d). A weak, non-specific signal of the peptide cross-linked to BSA was observed but only at the highest concentration where the peptide was in 2X molar excess to JNK1 (Fig. 2d). At this concentration, most of the peptide in solution is expected to be unbound to JNK1, which binds MKK4tides with a K_d_ of 3.5μM^5^ Thus, MKK4tide’s crosslinking to JNK1 likely reflects a domain-dependent binding event.

### LiF-MS reveals MKK4’s docking site on JNK1 and p38α

We next mapped the interaction site of MKK4tide on purified JNK1 using peptide I (Fig. 2a) and the LiF-MS workflow described in Fig. 1b. The resulting MS/MS data was analyzed to identify human proteins containing the 72-dalton butanol “mark” using COMET^17,18^. This revealed a set of Peptide Spectral Matches (PSMs) matching JNK1, a subset of which are shown as an example in Fig. 3a. We scored this data, removing low-scoring butanol sites (Supplementary Figure 1, see methods for detail). Since a structure of JNK1 complexed with MKK4 is not available, we mapped the location of each butanol-marked residue onto the published JNK1-NFAT4 structure (PDB 2XS0)^5^, comparing the location of the marked residues to the NFAT4 binding site (Supplementary Figure 2). While peptides both proximal to and distal from the docking site were discovered, the highest-scoring peptides mapped to the negatively charged common docking (CD) groove close to the putative N-terminus of MKK4 (Fig. 3b-c; MKK4 N-terminus, yellow; marked residue sidechains, red). Notably, we also observed high-scoring peptides in a loop distinct from the CD groove, although still potentially within range of an interaction with the flexible N-terminus of the peptide (Fig. 3d). We hypothesize this loop may be mobile in solution and could be a second part of the binding site that contacts the peptide. Together these data demonstrate the capabilities of LiF-MS to both probe the binding location and motion of linear motifs.

**Figure 3.**
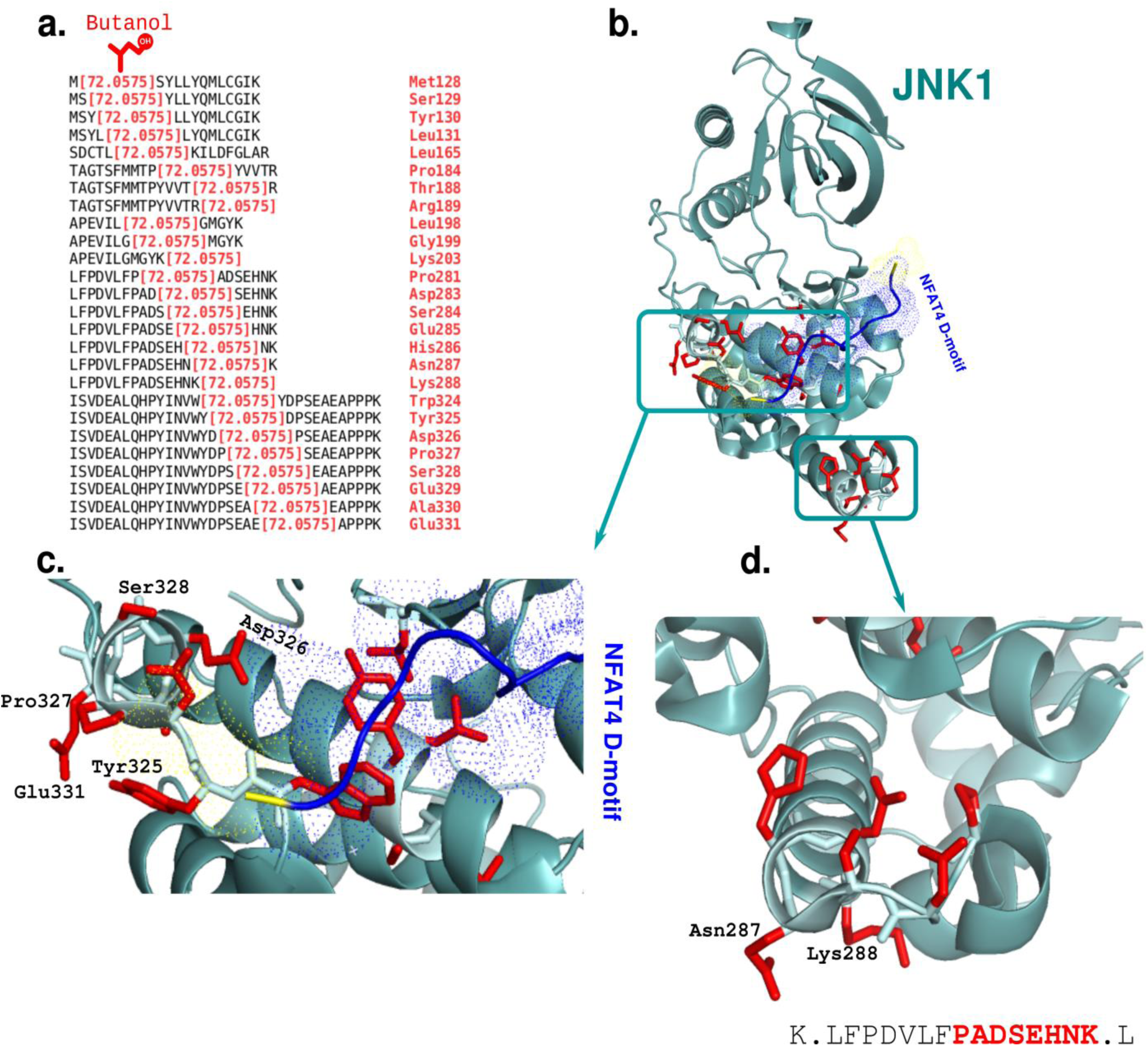
MKK4’s interaction with JNK1 mapped by LiF-MS. (a) Example list of butanol marked tryptic peptides derived from the crosslinking reaction. MKK4 peptide I was incubated with purified JNK1 isoform 3 and analyzed by LC-MS. (b-d) Peptides marked with butanol were mapped onto the JNK1-pepNFAT4 crystal structure (PDB ID: 2XS0). Sidechains of footprinted residues are shown in red as sticks. The MKK6 D-motif from the p38-MKK6 structure (PDB ID: 5ETF) is shown in blue. (b) Overview of the JNK1 structure with key footprinted regions highlighted. (c) Footprinted residues close to the CD-groove visible in the crystal structure. (d) Placement of a footprinted loop not proximal to the rest of the D-site suggesting extended conformational dynamics.

We then tested footprinting of MKK4tide on a related kinase, p38α, comparing it to JNK1 (Fig. 4a-b). Previous studies suggest MKK4 binds to the divergent kinases JNK1 and p38α/ERK2 docking sites though distinct conformations^5^. In fact, MKK4 is predicted to bind in a similar manner to members of the MKK6 family in complex with p38α^6^. Therefore, we overlaid our footprinting data from peptide I onto the p38α-MKK6 structure (PDB: 5ETF)^19^ (Fig. 4b). Similar to JNK1, we find the majority of marked residues to be proximal to the predicted binding site (Figure 4b). Interestingly, footprinting in this region suggests the MKK4tide “loops around” towards its C-terminus (Fig. 4b, cartoons) which is consistent with the predicted structure of p38α interacting peptides in the MKK6 family^6^. We found peptide I labeled a different region on p38α compared to JNK1, consistent with evidence that MKK4 binds in a different binding mode to JNK1 and p38a^5^ (compare Fig 4a and b).

**Figure 4.**
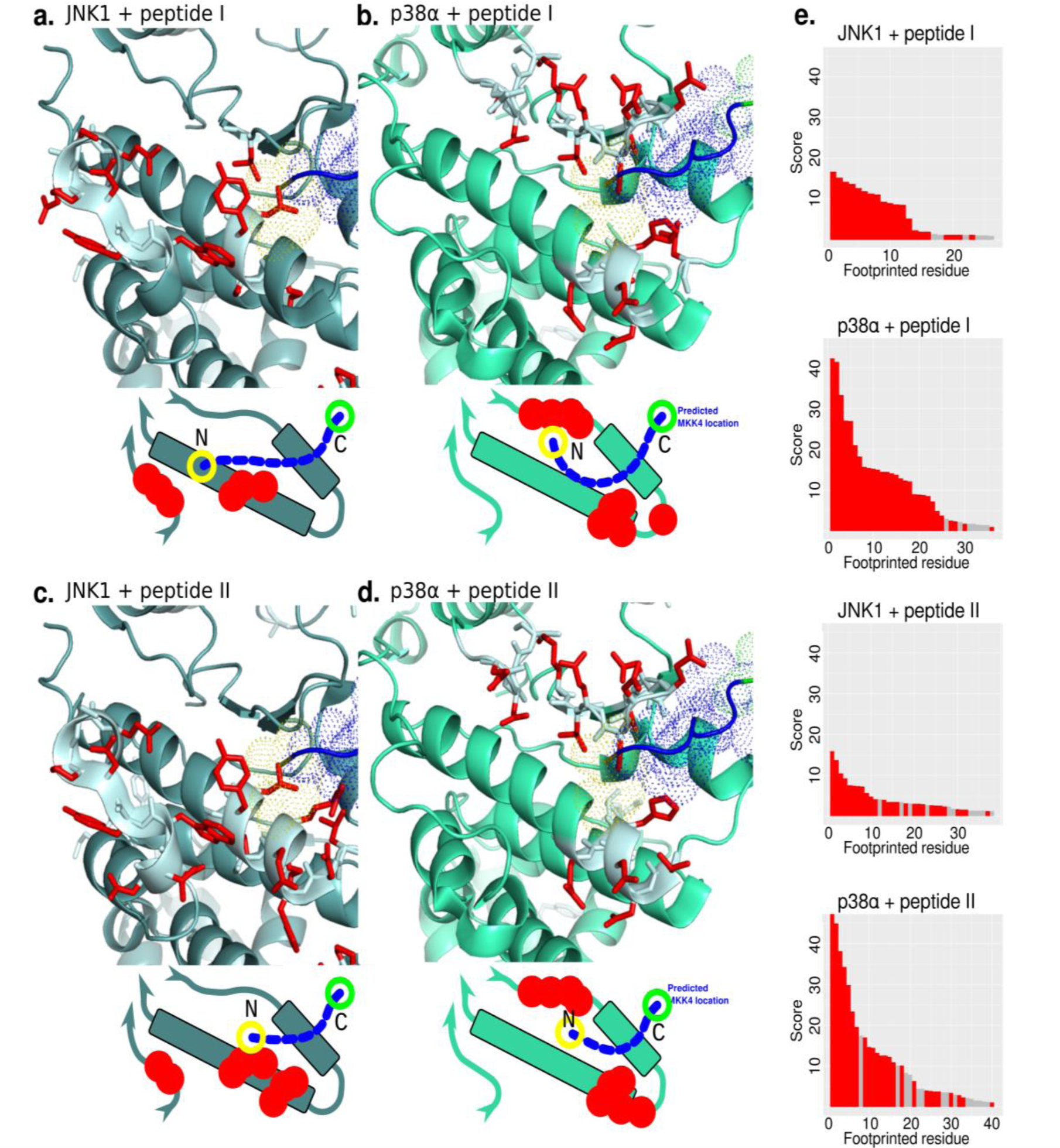
JNK1 and p38α’s docking site mapped with LiF-MS and compared. (a,b) MKK4 labels JNK1 and p38α at different faces of the D-site. Residues marked by crosslinker-containing peptide I were found at different helices in the acidic CD-groove of the MAPKs. (c,d) The CD-groove of JNK1 and p38α mapped using the shorter peptide II. The MKK6 D-motif from the p38α-MKK6 structure (PDB ID: 5ETF) is shown in blue for reference. Cartoons represent the predicted structure of the D-site. (e) Total butanol-labeled residues in each dataset were sorted in order of decreasing score, colored based on proximity to the MAPK D-site, and graphed. Red, crosslinking is close to wild-type D-site; grey, non-D-site-located crosslink.

Interestingly, we also found a region in p38α absent from the crystal structure that was marked by peptide I. Residues which extend above p38α’s docking site are labeled in our dataset (Supplementary Figure 3). Despite its distance from the docking site in the structure, we believe this strand is more flexible in solution and may form a disordered “lid” over bound D-motifs. Thus, we suggest LiF-MS can detect disordered interaction regions of binding sites not sufficiently ordered to be observed in a crystal structure. Overall, we show LiF-MS can be used to both form new structural hypothesis and confirm existing ones, especially in cases where other data is not available or not easy to obtain.

### Crosslinker placement can provide additional footprinting information

As linear motifs have much smaller binding surfaces than folded domains, peptide-protein interactions are much more prone to disruption by even small changes including point mutations, modified amino acids, and crosslinker placement^20^. To determine the effect of peptide length and crosslinker placement on our footprinting data, we synthesized MKK4 peptide II (Fig. 2a) in which the core D-motif-crosslinker distance was shortened by one amino acid. This peptide successfully crosslinked to all three kinases studied (Fig. 2b,c).

We then mapped the binding of peptide II onto JNK1 and p38α (Fig. 4c,d), comparing the results with peptide I (Fig. 4a,b). We then plotted both datasets by sorting butanol sites in order of decreasing score (Fig. 4e). We found the shorter MKK4tide resulted in a slight increase in the amount of crosslinked-residues that map outside of the predicted binding site (grey bars) when compared with marks at or near the predicted D-site docking cleft (red bars) (Fig. 4e, compare bottom to top). Furthermore, residues 123-127 in JNK1 were labeled with peptide II but not peptide I. This region is closer to the C-terminus of the peptide docking site consistent with a shortened N-terminal crosslinker placement (Fig. 4c, compare to Fig. 4a). Thus, for JNK1, the shorter peptide II revealed additional information about the MKK4-JNK1 binding surface. Mapping the binding of peptide II onto the p38α structure did not yield additional binding site information, though a slight increase in non-specific crosslinking was observed (Fig. 4e).

In conclusion, we find variation of the MKK4tide length and crosslinker placement affects footprinting data in kinase-specific ways. While shortening the peptide revealed a binding region on JNK1 not seen for the longer peptide (Compare Fig. 4a), the same experiment with p38α only created a noisier dataset (Fig. 4e). Thus, it is important for the user of this technology to vary peptide length and/or crosslinker placement in order to achieve optimal data.

### LiF-MS reveals MKK4 binds bi-directionally to ERK2

The MAP kinase ERK2, despite not being a known physiological target of MKK4, has been show to bind the MKK4 D-motif at its conserved docking site^5,8,16^. We found this intriguing, since ERK2 is not a physiological target of MKK4. Therefore, sought to footprint the binding of peptides I and II to ERK2’s docking site using LiF-MS. Peptide I yielded marked regions similar to both JNK1 and p38α, marking residues near ERK2’s docking site as expected (Fig. 5a, compare with Fig. 4). Surprisingly, we found that a region on the kinase located on the opposite end of the D-site was also marked extensively (Fig 5a,b, “reverse site”, compare purple and red dots on cartoons, bottom, also see Supplementary Figure 4). The reverse site was also marked by the shorter peptide II (Fig. 5b). Butanol marks in this region were consistent with the MKK4 D-motif binding in the opposite (C→N) orientation (Supplementary Figure 4, see cartoon). This region was scored highly, distributed throughout the dataset, and found in both peptide I and II experiments (Fig. 5c, compare red and purple).

**Figure 5.**
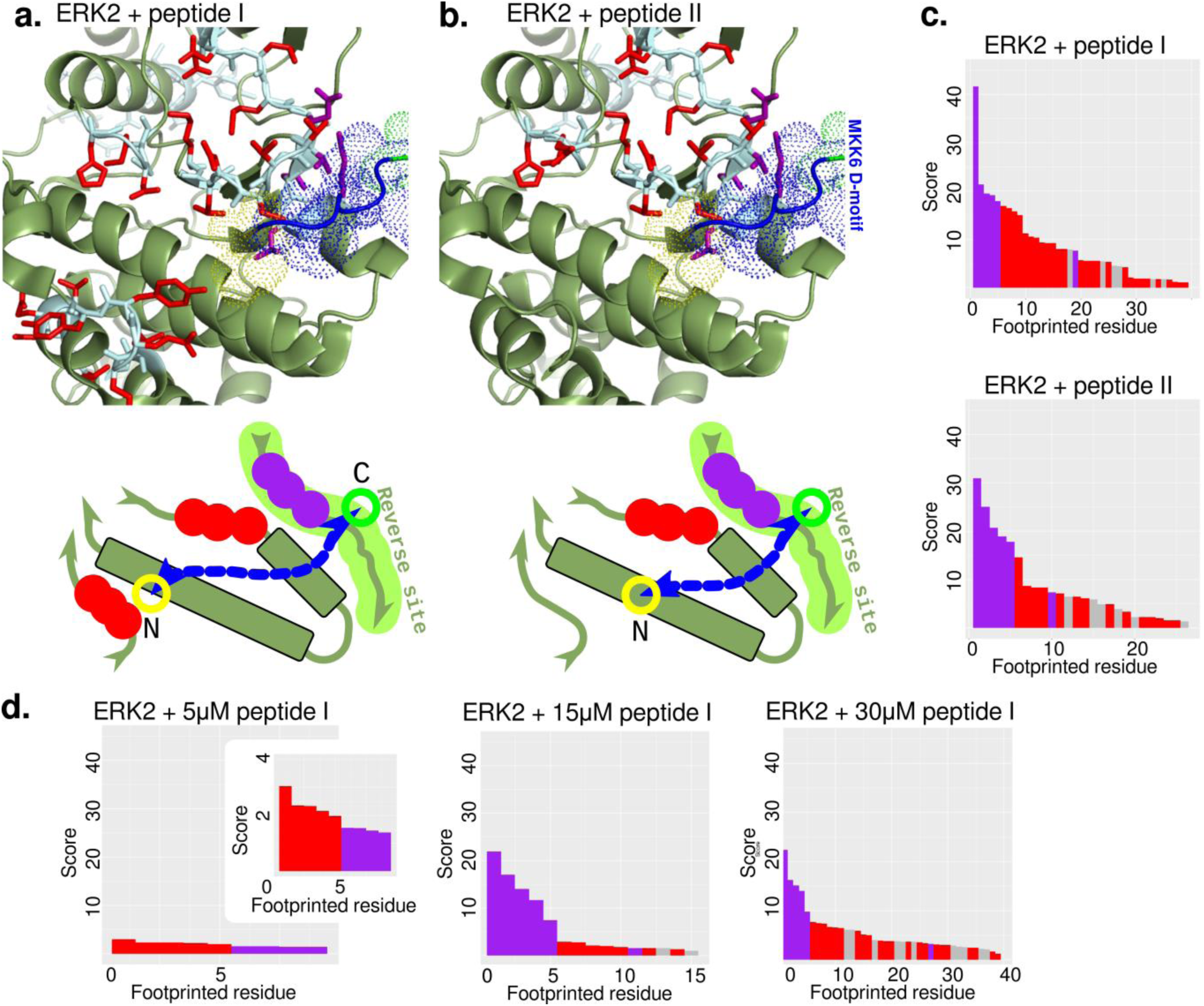
LiF-MS reveals bi-directional binding of MKK4 to ERK2. (a,b) MKK4 peptide I and II docking to ERK2 was mapped using LiF-MS. Red residues represent the known D-site; purple residues represent the discovered “reverse site.” Cartoons represent the predicted structure of the D-site. (c) Graph of butanol-modified residues sorted in order of decreasing score. Red, wild-type D-site; purple, reverse site; grey, non-D-site-located crosslinking. (d) Effect of increasing peptide concentration on detection of the reverse site. ERK2 was crosslinked to MKK4 peptide I at the indicated concentrations. Butanol-labeled residues were sorted in order by decreasing score. Scoring methodology same as in (c).

As relatively high concentrations of MKK4 D-motif and ERK2 were used in this experiment relative to the interaction affinity, we hypothesized that the marked reverse site regions would be a result of non-specific interactions saturating the ERK2 docking site. We performed LiF-MS with purified ERK2 and increasing concentrations of peptide I (Fig. 5d). Again, marked residues on both the predicted and the reverse sites were observed at all concentrations, though forward-binding regions had slightly higher scores at lower concentrations (Fig. 5d).

Overall, these ERK2 experiments establish LiF-MS as a platform to study the effects of varying parameters such as peptide length and concentration on a potentially difficult to study peptide-protein interaction. While our data suggests that MKK4 may be binding to the ERK2 docking site bi-directionally, additional work is needed to understand the biochemical relevance of this observation.

### Footprinting data assists in molecular docking

We hypothesized that butanol sites discovered by LiF-MS can reduce the computational search space when used as an anchoring point during molecular docking (Fig. 6a). We tested this using the docking program CABS-dock^21^, constraining a modified MKK4tide peptide I sequence to all identified butanol sites in the JNK1 D-site (Fig. 6a) to a non-peptide-bound JNK1 structure^21,22^. We scored backbone-only RMSDs of resulting structures against a modeled JNK1-MKK4 D-motif structure which uses NFAT4 as a structural template. Each CABS-dock run generates 10 top-ranked peptide structures from the initial set based on *k*-medoid clustering, and we took the best structure out of three separate runs as described previously^21^.

**Figure 6.**
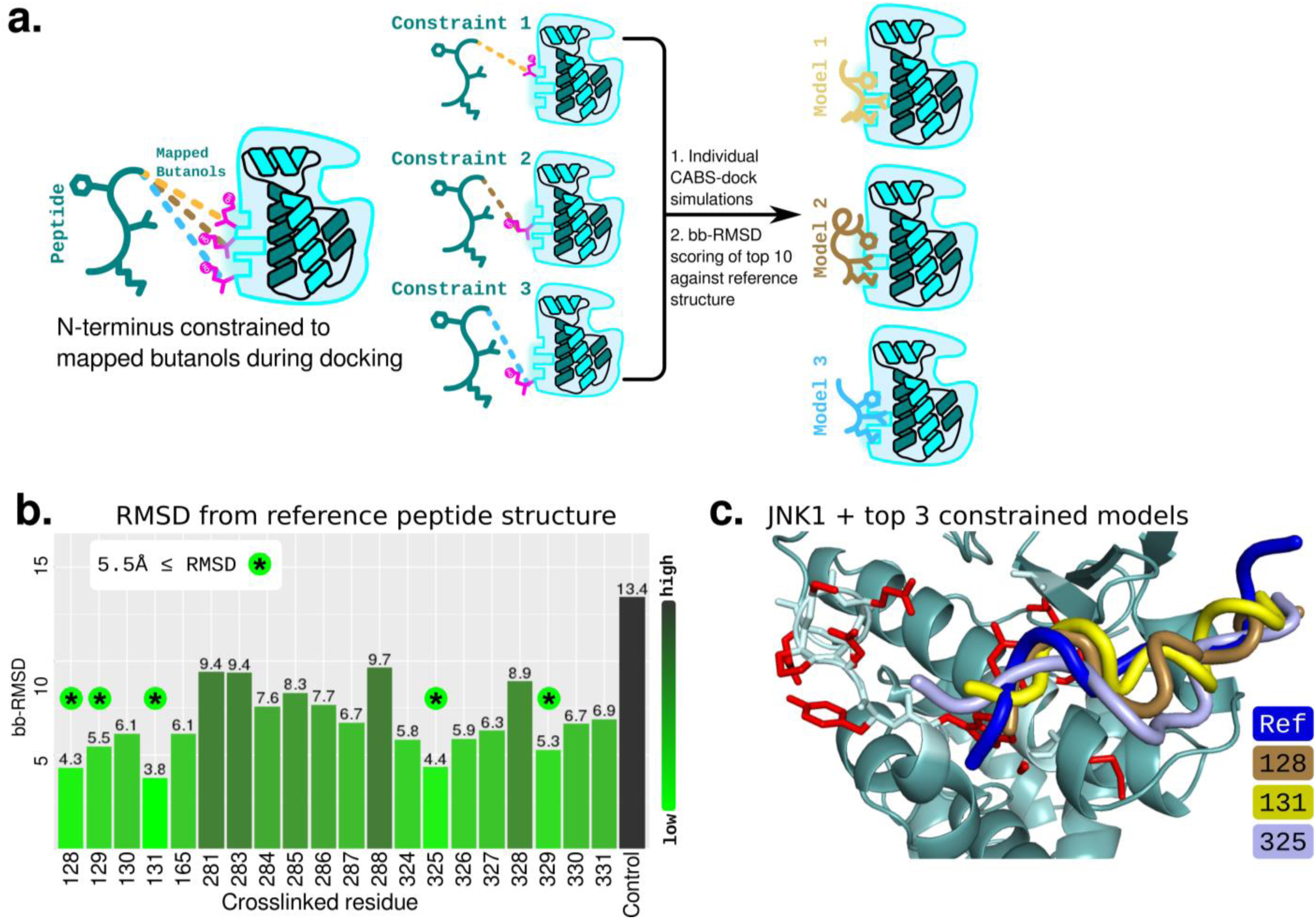
LiF-MS generated constraints assist in molecular docking. (a) Overview of modeling procedure. Docking peptide is constrained individually to each butanol-marked residue in the D-site and docked with CABS-dock along with an unconstrained control. The top 10 peptide structures are scored against a reference. (b) The MKK4 D-motif was docked to JNK1 (PDB: 3PZE) constrained to the indicated amino acid residues alongside an unconstrained experiment (Control). Independent CABS-dock runs were completed for each individual residue. Models in the top 10 were scored by backbone RMSD against a reference and the highest quality models from 3 independent runs compiled. Models with sufficient quality for near-native modeling are marked with a star (^*^). (c) Highest quality results overlaid on the reference (Ref, blue) MKK4 D-motif model used for scoring. Peptides in each experiment were constrained to the JNK1 amino acid numbers indicated.

We find that constraining MKK4 to known butanol sites produces consistently higher quality models than using unconstrained sequences alone in the top 10 models (Fig. 6b). We note that in all cases, the lowest backbone-RMSD found in the top 10 from butanol-constrained simulations was higher quality than the unconstrained control. Out of the 20 marked D-site residues used as constraints, five produced top model sets with an RMSD of ≤ 5.5Å, sufficient for near-native modeling by applications such as PepFlexDock^23^. Intriguingly, repeating this technique using a JNK1 structure in a D-motif bound conformation (PDB: 2XS0) gave lower model quality in our hands (Supplementary Figure 5a,b). We further found that anchoring the peptide to JNK1 residues 128, 131, and 325 gave the best models when compared with the reference control (Fig. 6c).

When analyzing the full trajectories (all 10,000 models per replicate, 30,000 models total per butanol site), gains in RMSD were also observed when compared with the unconstrained control. Importantly, constraining MKK4tide to several butanol sites in both bound and unbound JNK1 structures produced models with RMSD < 3Å considered to be “high” quality by the CABS-dock scoring prediction^21^, where controls were ≥ 3Å (Supplementary Figure 5c). The improvement in model quality with constrained peptides is noteworthy given that D-motifs in general are difficult to dock using CABS-dock, as was found in the case of the Ste7 D-motif to Fus3^24^.

## Discussion

Chemical crosslinking has been used extensively to probe the makeup and detailed structure of protein complexes^25^. Traditionally, structural data has been derived by crosslinking purified protein complexes with relatively non-specific crosslinkers such as disuccinimidyl suberate (DSS) or 1-ethyl-3-(-3-dimethylaminopropyl) carbodiimide hydrochloride (EDC), which link residues such as lysine and glutamate. Furthermore, most work has tried to elucidate the structure of folded protein-protein interactions, rather than map intrinsically disordered proteins. Notably, the structure of the intrinsically disordered “guardian of the genome,” p53, was probed using traditional homobifunctional crosslinkers^26,27^. Probing these unstructured interactions is considered one of the future targets of structural Mass Spectrometry^28^.

Here, we introduce LiF-MS, a novel method for footprinting the binding interface between disordered peptides and protein domains. This technique offers unique advantages and is a novel addition to current chemical crosslinking-mass spectrometry (XL-MS) methods.

First, the sequence simplicity of SLiMs, which form synthetic bait peptides ~10-15aa, allows crosslinker attachment through an endogenously introduced cysteine. This permits the synthesis yield and efficiency of a chemical crosslinker while maintaining the site-specificity of a genetically encoded crosslinker. This greatly simplifies synthesis of the bait SLiMs, permitting production of the high concentrations necessary to map weak, transient interactions.

Second, the crosslinker’s acid cleavability permits a straightforward enrichment and elution strategy which concentrates crosslinked material after trypsiniziation. Only a small fraction of peptide actually covalently bonds to the folded protein due to the inefficiency of the diazirine warhead; as biotinylated SLiMs can be used to isolate crosslinked peptides from a large pool of trypsinized material, this signal can be easily amplified. We believe this is crucial for mapping these type of interactions, which are weak, transient, and mobile.

Concurrent to this work, a cleavable, photoactivatable diazirine-based crosslinker has been used to map the conformational changes of the acid chaperone HdeA^29^. Cleavage of this crosslinker with hydrogen peroxide leaves a residual mark which can be modified with dyes or other functional groups ^30,31^. Mapping protein-protein interactions with this reagent has been utilized successfully. While the butanol group produced in LiF-MS cannot be modified, we believe the ease of synthesis of large amounts of prey peptide through cysteine alkylation (Fig. 1a) rather than codon suppression gives LiF-MS an advantage when specifically mapping transient interactions in a purified context.

Furthermore, LiF-MS can provide crucial and complementary information to currently utilized structural techniques. In particular, the site-specific footprinting maps produced by LiF-MS may be combined with NMR data to produce a more accurate picture of how an unstructured region contacts a folded domain. Also, elements not visible in X-ray crystal structures can be detected with LiF-MS as in the case of the MKK4tide-p38α interaction (Supplementary Figure 3). LiF-MS may further be used in tandem with H/D exchange or other footprinting techniques to verify the broad location of a disordered interaction, as both methods can probe the effect of conformational dynamics on protein-protein binding sites. Additionally, an entire binding site could be mapped through systematic placement of the reactive cysteine along the length of the peptide.

As LiF-MS detects the motion of a single labeled “point” on the peptide studied, it can uniquely determine the binding conformation and directionality of a disordered peptide within a protein domain. We demonstrate MKK4 binds JNK1 and p38α in divergent conformations as predicted previously, confirming a previous structural hypothesis^5,6^. Interestingly, a majority of MKK6’s basic N-terminus is invisible in the p38α-MKK6 crystal structure, suggesting intrinsic disorder which may be similar for the MKK4 interaction^19^.

MKK4’s seemingly bi-directional interaction with ERK2 is particularly curious. The ERK2 D-site may be unusually promiscuous and bind peptides regardless of sequence, though ERK2 contains other docking sites for motifs such as FXFP and a catalytic cleft which were not labeled significantly; we therefore believe this is unlikely. Regardless, this type of directionality data would be difficult to obtain using other biophysical methods and may offer unique insights into the structure of peptide-protein complexes.

We demonstrate here a novel method for probing the footprint of an unstructured peptide on a folded target protein using mass spectrometry. This technique provides amino-acid level interaction data and can be used to elucidate peptide dynamics within a folded domain. In summary, LiF-MS complements other *in vitro* XL-MS technologies and offers a robust method for studying the interactions of IDPs.

## Methods

### Cloning MAP kinases

JNK1 β-1 isoform lacking 20 amino acids from the C-terminus (JNK1 ΔC20), ERK2, and p38α α-isoform were cloned into pBH4 as described. All constructs were transformed into BL21 (DE3) RIL cells and overexpressed in Terrific Broth for 4 hours at 25°C with 0.2mM IPTG.

### Purification of JNK1 kinase

Cells were pelleted and resuspended in lysis buffer (20mM Tris, 150mM NaCl, 10% Glycerol, pH 8.0), incubated with lysozyme, and lysed at 20 000psi with an Emulsiflex homogenizer. Lysate was loaded onto 0.5mL Ni-NTA agarose (Qiagen) and washed with 20 column volumes (CV) of lysis buffer + 20mM imidazole. Protein was eluted with lysis buffer + 250mM imidazole, combined with 1/100 (OD_280_/OD_280_) of purified TEV protease, and dialyzed into ion exchange buffer (20mM Tris, 20mM NaCl, 10% Glycerol, 20mM DTT, pH 8.0) overnight. Dialyzed protein was loaded onto a hand-poured column of 5mL Q Sepharose Fast Flow resin (Amersham Biosciences). After washing with 10CV of ion exchange buffer, protein was eluted using a gradient to 1M NaCl. Final fractions were concentrated to 15mg/mL, flash frozen in liquid nitrogen, and stored at -80°C.

### Purification of ERK2 and p38α kinases

Cells suspended in lysis buffer (50mM Tris, 150mM NaCl, 10% Glycerol, pH 8.0) were incubated with 1mg/mL lysozyme at 4°C and lysed with 0.1% (w/v) sodium deoxycholate. Nucleic acid was precipitated with 0.1% (w/v) branched polyethyleneimine (Sigma). Supernatant was incubated with Ni-NTA agarose. Resin was washed with 10CV lysis buffer, 10CV lysis buffer + 20mM imidazole, and eluted with lysis buffer + 100mM imidazole. Resulting protein was cleaved with TEV protease and dialyzed into dialysis buffer (50mM Tris, 150mM NaCl, 10% Glycerol, 2mM DTT). Protein was further purified with subtractive purification using Ni-NTA agarose. Fractions were diluted in 100% glycerol to a final concentration of 55%, flash frozen in liquid nitrogen, and stored at -80°C.

### Synthesis of D-motifs

The MKK4 base D-motif with sequence QGKRKALKLNFAN was used as described^5,8^. Peptide wt (wild-type) was ordered with the sequence CQGKRKALKLNFANW. The shorter peptide I was synthesized with sequence CGKRKALKLNFANW. Peptide II was synthesized with sequence CKRKALKLNFANW. Peptides wt, I, and II were ordered from Genscript, contained an N-terminal biotin, and were >85% pure. The C-terminal tryptophan was included to permit spectroscopic concentration measurements.

### Synthesis of crosslinker

Synthesis of the crosslinker was performed as described^11^.

### Alkylation of peptides

Crosslinker is conjugated to peptide cysteines through an iodoacetamide moiety. Peptides were dissolved in crosslinking buffer (CB; 50mM Tris pH 8.3, 150mM NaCl) and diluted into CB to a final concentration of 200-500μM. 0.5M crosslinker in DMF was added to a final concentration of 2.5mM. Samples were incubated at 4°C for 90 minutes in 1.5mL black centrifuge tubes (Argos Technologies, Inc.) with gentle nutation. Excess crosslinker was then destroyed by the addition of DTT to 5mM. Alkylation efficiency was assayed by MALDI-TOF mass spectrometry.

### Crosslinking of kinases

All steps were performed at 4°C. Purified MAP kinases were exchanged into CB using hand-poured 1mL spin columns containing Sephadex G-25 (Sigma). Alkylated peptide was added to purified kinase and incubated in black centrifuge tubes for 30 minutes with over-end nutation and irradiated with a 4-watt UVL-21 BLAK-RAY^®^ 365nm ultraviolet lamp in a clear 96-well V-bottom storage plate (Corning, Inc) at a distance of 1-2cm from sample. Lamp/plate assembly were covered in aluminum foil to increase sample photoyield. Reagent amounts and concentrations in each experiment are described in Supplementary Table 1.

### Trypsinization and mass spectrometry

Crosslinked MAP kinase was incubated with 5mM DTT for 45 minutes at 37°C and 15mM iodoacetamide for 20 minutes at room temperature in the dark. 1 μg of Trypsin Gold (Promega) was added and sample incubated at 37°C overnight with gentle nutation. Digested sample was spun down to remove precipitated peptides and soluble fraction kept. 10μL high-capacity streptavidin agarose (Thermo) in CB was added and the sample nutated at 4°C for 1 hour. Resin was washed 3X with 4mM KH_2_PO_4_, 16mM Na_2_HPO_4_, 115mM NaCl, 5% acetonitrile, pH 7.3 and 2X with double distilled water as described^32^. Peptides were eluted by heating in 4% (v/v) formic acid at 55°C for 2 hours, desalted with reverse-phase C18 spin columns, and analyzed by LC-MS/MS on a Q Exactive HF Orbitrap (Thermo).

### Western blots

Kinases were separated in 12% SDS-PAGE gels and transferred to Immobilon-FL membrane (Millipore), blocked with TBS Odyssey Blocking Buffer (Li-Cor), and probed with anti-biotin antibody (Santa Cruz Biotechnologies, Inc) or IRDye^®^ 800CW Streptavidinin (Li-Cor) in TBS-T + 10% Odyssey Blocking Buffer. Blots were imaged on a Li-Cor Odyssey Clx.

### Proteomics

#### Peptide identification using MS/MS spectra

See Supplementary Figure 1. MS/MS spectra were searched using MS/MS database search tool Comet 2017.01 rev. 2^17,18^ against the SwissProt human protein sequence database modified to contain the JNK1 β-1 isoform. The following parameters were used: precursor mass tolerance of 20 ppm, tryptic peptides with up to 3 missed cleavages, allowing a butanol modification (+72.0575 mass shift) on any amino acid as well as oxidized Met (+15.9949) and deamidations of Gln and Asp (+1Da) as variable modifications. PSM lists were filtered (see next session) using the R statistical computing environment (version 3.3.3)^33^ with dplyr version 0.7.4^34^ and stringr 1.2.0^35^ packages; the ggplot2 package was used to plot spectra^36^.

#### Butanol scoring and mapping

PSMs not from the protein being studied, lacking the butanol modification, or with a COMET expectation value (E-value) greater than 0.1 were removed. E-values were converted to NLEV (Negative Log E-value, -log_10_[E-value]). Non-butanol modifications were removed from the dataset and duplicate PSMs grouped, summing the NLEVs of all supporting PSMs to compute a final “score” for each modification site (Supplementary Figure 1). The location of each “site” with a butanol was established from the protein sequence and amino acid residues containing the modification were mapped using pymol^37^. The final dataset contains all butanol-modified amino acid residues identified by at least one PSM with E-value below 0.1 cutoff (NLEV score above 1).

#### Peptide docking

CABS-dock standalone version 0.8.8 was used for docking experiments^21^. MKK4 peptide I (GKRKALKLNFAN) was constrained to butanol sites derived from the crosslinking experiment with a distance constant of 12.0 and weight of 1.0. All residues labeled near the known D-site were used as docking constraints. Crystal structures from peptide-bound (PDB: 2XS0)^5^ and unbound (PDB: 3PZE)^22^ JNK1 were used as folded receptors.

#### Scoring of output structures

As no MKK4 D-motif crystal structure was available, the NFAT4 D-motif crystal structure (PDB: 2XS0)^5^ was used as a template. The NFAT4 D-motif was mutated to match the MKK4 peptide I sequence (GKRKALKLNFAN) based on sequence alignment using pymol and the resulting synthetic “reference” MKK4 peptide structure refined using Rosetta PepFlexDock^23^. RMSDs of CABS-dock models from the reference MKK4 peptide were calculated using the bio3d R package^38^.

## Acknowledgements

We thank Felipe da Veiga Leprevost for help with MS/MS data analysis. We thank Gergő Gógl and Reményi Attila for contributing the pBH4 vectors containing the JNK1, ERK2, and p38α sequences. We thank Jennifer Brace for critical review and editing of the manuscript. This work was funded in part by grants from the NIH (R01GM94231 and U24CA210967 to A.I.N.).

Proteomics services were performed by the Northwestern Proteomics Core Facility, generously supported by NCI CCSG P30 CA060553 awarded to the Robert H Lurie Comprehensive Cancer Center and the National Resource for Translational and Developmental Proteomics supported by P41 GM108569.

This work used resources of the Northwestern University Structural Biology Facility, which is generously supported by NCI CCSG P30 CA060553 awarded to the Robert H. Lurie Comprehensive Cancer Center.

## Author contributions

B.P. designed, optimized and performed the crosslinking methodology. E.G. did preliminary protein biochemistry and crosslinking experiments. B.P. and A.N. performed mass spectrometry data processing and scoring. A.S. and D.K. provided reagents and assistance with synthesis of the crosslinker. B.P. did computational modeling and wrote the paper. E.L.W conceived and supervised the project.

